# POT-3 preferentially binds the terminal DNA-repeat on the telomeric G-overhang

**DOI:** 10.1101/2022.07.01.497317

**Authors:** Xupeng Yu, Sean Gray, Helder Ferreira

**Author notes:** Equal contribution (Xupeng Yu, Sean Gray).

## Abstract

Eukaryotic chromosomes typically end in 3’ telomeric overhangs. The safeguarding of telomeric single-stranded DNA overhangs is carried out by factors related to the Protection of Telomeres 1 (POT1) protein in humans. Of the three POT1-like proteins in *C. elegans*, POT-3 was the only member thought to not play a role at telomeres. Here, we provide evidence that POT-3 is a *bona fide* telomere-binding protein. Using a new loss-of-function mutant, we show that the absence of POT-3 causes telomere lengthening and increased levels of telomeric C-circles. We find that POT-3 directly binds the telomeric G-strand *in vitro* and map its minimal DNA binding site to the six-nucleotide motif, GCTTAG. We further show that the closely related POT-2 protein binds the same motif, but that POT-3 shows higher sequence selectivity. Crucially, in contrast to POT-2, POT-3 prefers binding sites immediately adjacent to the 3’ end of DNA. These differences are significant as genetic analyses reveal that *pot-2* and *pot-3* do not function redundantly with each other *in vivo*. Our work highlights the rapid evolution and specialisation of telomere binding proteins and places POT-3 in a unique position to influence activities that control telomere length.

## Introduction

Telomeres are large protein-DNA structures that protect the ends of linear chromosomes from inappropriate DNA repair and the end replication problem. Importantly, the protective functions of telomeres are mediated by proteins rather than the underlying DNA sequence *per se* [1]. These protective, telomere-associated proteins tend to form a complex *in vivo*, best exemplified by the human Shelterin complex [2]. Single-stranded DNA (ssDNA) binding proteins are important components of the Shelterin complex as, in most species, telomeres are processed to form a 3’ ssDNA overhang [3]. These overhangs help to distinguish telomeric DNA repeats at chromosome ends (true telomeres) from internal repeats of interstitial telomere sequences (ITSs). In humans, this telomeric ssDNA binding function is carried out by the POT1 (Protection of Telomeres 1) protein [2]. The telomere protective functions of POT1 is highly dependent on its ssDNA binding specificity through a conserved oligosaccharide/oligonucleotide binding fold (OB-fold) [4, 5].

The composition of OB folds is defined by five antiparallel β strands forming a distinct β barrel [4]. Variable loops connect these secondary structure elements and have a significant role in forming the binding site [6]. They are also primarily responsible for variability in OB-fold lengths between 70-150 amino acids [7]. OB folds are present across evolutionarily distant organisms and their ligands can range from RNA and ssDNA to protein [7], although they most commonly bind ssDNA. Indeed, OB fold proteins play crucial roles in processes such telomerase activity that generate ssDNA.

Within human Shelterin, POT1 is the only protein that confers ssDNA binding. It binds telomeric DNA via two OB folds and deletion of POT1 results in telomere-associated dysfunctions such as 5’ end resection, and telomere elongation [8]. Mice and rats have two POT1 proteins, referred to as mPOT1a and mPOT1b [9]. These are homologous to the two OB folds of human POT1 [10]. Despite having duplicated only recently and displaying 75% sequence similarity, the functions of mPOT1a and mPOT1b have diverged significantly [9].

The nematode worm *C. elegans* contains three POT1-like proteins (POT-1, POT-2 and POT-3), each containing a single OB fold. Loss of *C. elegans* POT-1 or POT-2 results in telomere elongation [11], mimicking what is seen in humans. The closely related *pot-3* gene shares very high sequence similarity to *pot-*2 (∼60% amino acid identity). However, POT-3 initially appeared to have no telomeric function, as a *pot-3(ok1530)* allele was shown to have normal telomere length [12]. Here, we characterise the behaviour of the POT-3 protein *in vitro* and the phenotypes of a novel *pot-3* mutation *in vivo* to show that POT-3 does indeed have an important telomeric function. Understanding OB-fold containing telomere binding proteins from different eukaryotes may shed light on the diversity of telomere maintenance mechanisms.

## Results

### Mutation of *pot-3* increases telomere length and recombination

Given the homology to POT-2, we suspected that POT-3 might also play a role at telomeres. To test whether mutation of *pot-3* caused a telomeric phenotype, we isolated a new null allele, *pot-3(syb2415)* containing a 500bp deletion spanning the entire OB fold (Figure 1A). *pot-3(syb2415)* worms are viable and fertile. Interestingly, we observed that *pot-3(syb2415)* mutants have markedly longer telomeres than wildtype worms 5.5 - 23.9 Kbp vs 2.2 - 8.6 Kbp respectively (Figure 1B and Supplementary Table 1). We observed that telomere length in *pot-3* mutants is not stable but rather increases over time (data not shown). This behaviour has previously been seen with *pot-2* mutants [12]. Therefore, to accurately compare telomere lengths, all experiments were carried out with early generations of the relevant genotypes.

**Figure 1.**
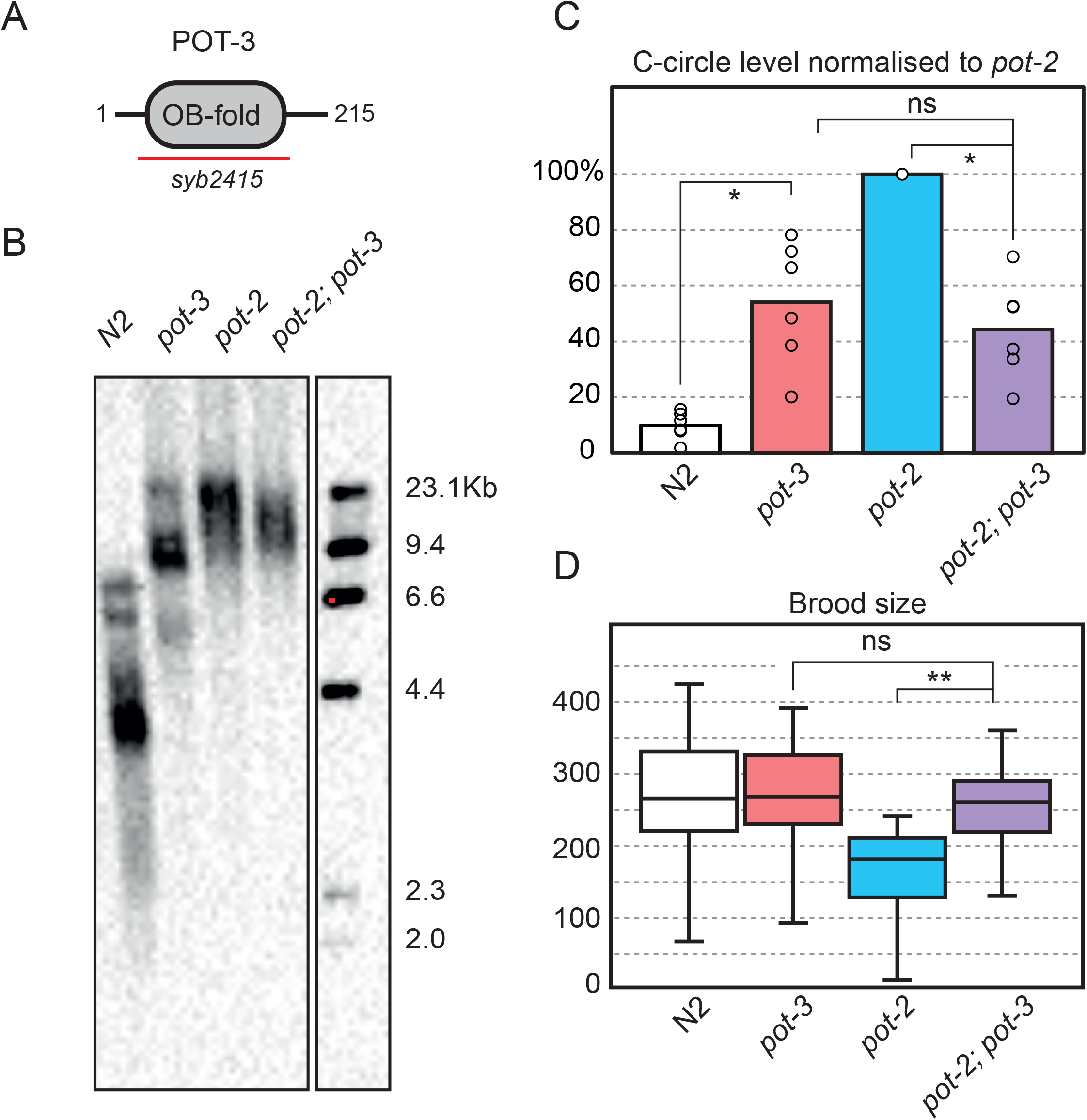
Mutation of *pot-3* increases telomere length and recombination in a manner that is epistatic with *pot-2*. **A**. POT-3 is 215aa long, the *syb2415* allele contains a 500bp deletion spanning the entire OB-fold region. **B**. Southern blot of terminal restriction fragments from genomic DNA show that mutation of *pot-3* results in an increase in telomere length almost to that of *pot-2*. Interestingly *pot-2; pot-3* double mutants do not have longer telomeres than either single mutant. Membrane was probed with a DIG-labelled (TTAGGC)_4_ oligo. **C**. Telomeric C-circle assays were carried out using phi29 polymerase, spotted onto a nitrocellulose membrane and probed with a DIG-labelled (TTAGGC)_4_ oligo. Signal intensity was quantified using ImageJ and results plotted relative to *pot-2* (set to 100%). The bar graph shows the average of six independent experiments with individual results displayed as open circles. Mutation of *pot-3* increases C-circle levels above that of wildtype but supressed the levels of C-circles in a *pot-2* background, ns = not significant, * p = <0.05 (T-test, two tailed distribution, unequal variance). **D**. The total number of viable offspring per adult worm (brood size) was measured for the indicated genotypes at 20°C. The Box plot displays the mean, 25th and 75th percentile from at least 20 independent adults. Mutation of *pot-3* has no significant effect on brood size on its own but supresses the lower brood size of a *pot-2* mutant. ns = not significant, ** p = <0.005

Besides telomere lengthening, *pot-2* mutants also display higher levels of telomeric C-circles [11]. These extra-chromosomal circles of telomeric DNA are hallmarks of the alternative lengthening of telomeres (ALT) pathway [13]. Strikingly, we also observe that *pot-3* mutants have significantly higher levels of C-circles than wildtype (Figure 1C). Thus, we see that *pot-3* mutants show both of *pot-2*’s telomere phenotypes, namely longer telomeres and increased levels of C-circles. These data strongly suggest that POT-3 plays a similar role to that of POT-2 at worm telomeres.

### POT-3 does not act redundantly with POT-2 *in vivo*

We noticed that, although similar to *pot-2*, the telomeric phenotypes of *pot-3* mutants were always weaker. This raised the possibility that perhaps POT-2 and POT-3 were carrying out the same role (*i*.*e*. they were redundant) but that POT-2 was somehow more important or abundant. If this were the case, we would observe stronger telomeric defects in a *pot-2; pot-3* double mutant compared to either single mutant. However, we found that a *pot-2; pot-3* double mutant had weaker telomeric phenotypes than a *pot-2* single mutant. The loss of *pot-3* did not exacerbate but rather supressed both the telomere length and telomere recombination phenotypes of *pot-2* mutants (Figure 1B, C and Supplementary Table 1). Thus, rather than acting redundantly, *pot-2* and *pot-3* mutants are epistatic. This indicates that they do not carry out the same function but work together within the same genetic pathway. Indeed, this epistatic relationship is not restricted to telomeric phenotypes. We find that *pot-2; pot-3* double mutants also have a significantly higher brood size than *pot-2* single mutants (Figure 1D). This indicates that *pot-2* and *pot-3* also affect general fitness in an epistatic manner.

Loss of POT homologs in yeasts leads to telomere uncapping, resulting in loss of telomeric DNA and chromosome fusion [14, 15]. Although *pot-2* single mutant worms did not show signs of chromosome fusion [16], we wondered whether this phenotype might only be revealed in a *pot-2; pot-3* double mutant. The number of chromosomes in *C. elegans* can readily be counted in meiotic cells in diakinesis. However, we did not find evidence of large-scale chromosome fusions in either the *pot-3* single or the *pot-2; pot-3* double mutant (data not shown).

### POT-3 specifically binds the G-strand of telomeric DNA

To understand how POT-3 might be acting on telomeres *in vivo*, we decided to test its DNA-binding properties *in vitro*. His-tagged POT-3 was recombinantly expressed in *E. coli* and purified to homogeneity using a combination of affinity chromatography and gel filtration (Figure 2A, B). Most POT homologs from other species contain multiple OB-folds. In contrast, all *C. elegans* POT homologs contain only a single OB fold. We therefore wondered whether they might multimerise. However, interestingly, POT-3 is monomeric in solution, although it is prone to forming cysteine-mediated dimers (Figure 2A, B).

**Figure 2.**
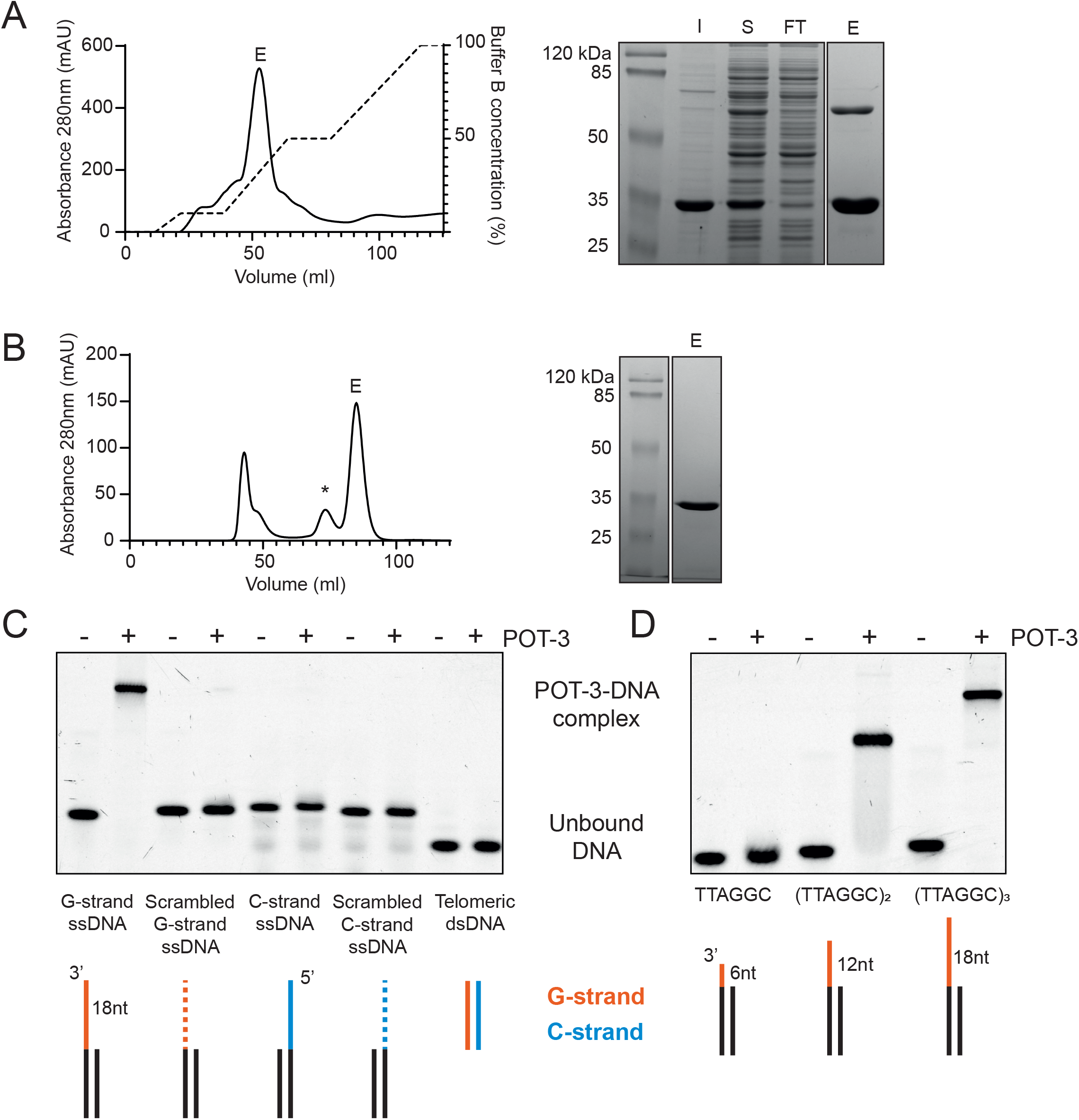
POT-3 is a monomer and binds the telomeric G-strand as ssDNA. **A**. Nickel affinity purification of 6xHis-tagged POT-3 expressed from *E. coli*. The panel on the right is a Coomassie stained SDS-PAGE gel of different fractions collected during the purification: I - insoluble, S - soluble, FT - flowthrough, E - elution. **B**. Affinity purified POT-3 was subsequently run over a HiLoad 16/60 Superdex 200 size exclusion column. The majority of POT-3 migrates as a monomer although it can form disulphide-mediated protein dimers, indicated by an asterisk. **C**. 500nM Purified POT-3 was incubated with 50nM Cy5-labelled DNA and run on a native acrylamide gel. DNA substrates contained a non-telomeric dsDNA region (black) with either a 3’ G-strand telomeric ssDNA overhang (orange) or a 5’ C-strand telomeric ssDNA overhang (blue). Telomeric ssDNA overhangs were 18 nucleotides long and consisted of three telomeric repeats, (TTAGGC)_3_. The scrambled G and C-strand overhands retained the same GC content as the telomeric sequence but the nucleotide order was randomised. **D**. POT-3 only binds to G-overhangs containing more than one copy of the telomeric sequence TTAGGC.

To assay POT-3’s specificity for telomeric DNA, we hybridised DNA oligonucleotides to generate structures that mimicked telomeres. These double-stranded molecules had single-stranded DNA overhangs containing the *C. elegans* telomere sequence. Using electrophoretic mobility shift assays (EMSA), we find that POT-3 efficiently binds DNA templates containing a G-overhang (Figure 2C). This interaction is DNA sequence-specific because if we scramble the G-overhang sequence then POT-3 no longer recognised it. Moreover, POT-3 binding is strand-specific. If we use templates containing the complementary telomeric C-overhang, POT-3 fails to bind. Recognition of the telomeric G-strand has to take place within the context of ssDNA because telomeric dsDNA is not bound by POT-3 (Figure 2C). These data are consistent with the observed telomeric phenotypes of *pot-3* mutant worms and strongly suggest that POT-3 is a *bona fide* telomere binding protein.

### POT-3 and POT-2 bind a minimal six nucleotide GCTTAG motif

We initially thought that POT-3 needed more than six nucleotides of telomeric G-strand to bind. Using templates containing either one, two or three repeats of the telomeric sequence (TTAGGC) as ssDNA, we saw that POT-3 would readily bind overhangs containing two or more copies of TTAGGC but could not bind an overhang with a single repeat (Figure 2D). However, it remained possible that that we were missing the minimal binding motif of POT-3 because we were not using the correct register of telomeric repeat sequence.

To address this, we tested POT-3 binding to a series of telomeric substrates. These started with a single six nucleotide TTAGGC overhang (which we knew POT-3 did not bind) that were extended one nucleotide at a time at the 3’ end until it reached a twelve nucleotide (TTAGGC)_2_ overhang (which we knew POT-3 did bind). We reasoned that if we identified the point at which extending the 3’ end resulted in POT-3 binding, we could then shorten the telomeric repeat from the 5’ end to identify the minimal binding motif. In contrast to what we previously thought, POT-3 does indeed only require six nucleotides for complete binding. However, this has to be in the register GCTTAG and not TTAGGC (Figure 3A). This is comparable to the binding behaviour of human POT1. Although, the two OB folds of human POT1 bind a ten nucleotide motif, this also ends in TTAG. Moreover, human POT1 has a higher affinity for (GGTTAG)_2_ than (TTAGGG)_2_ [17].

**Figure 3.**
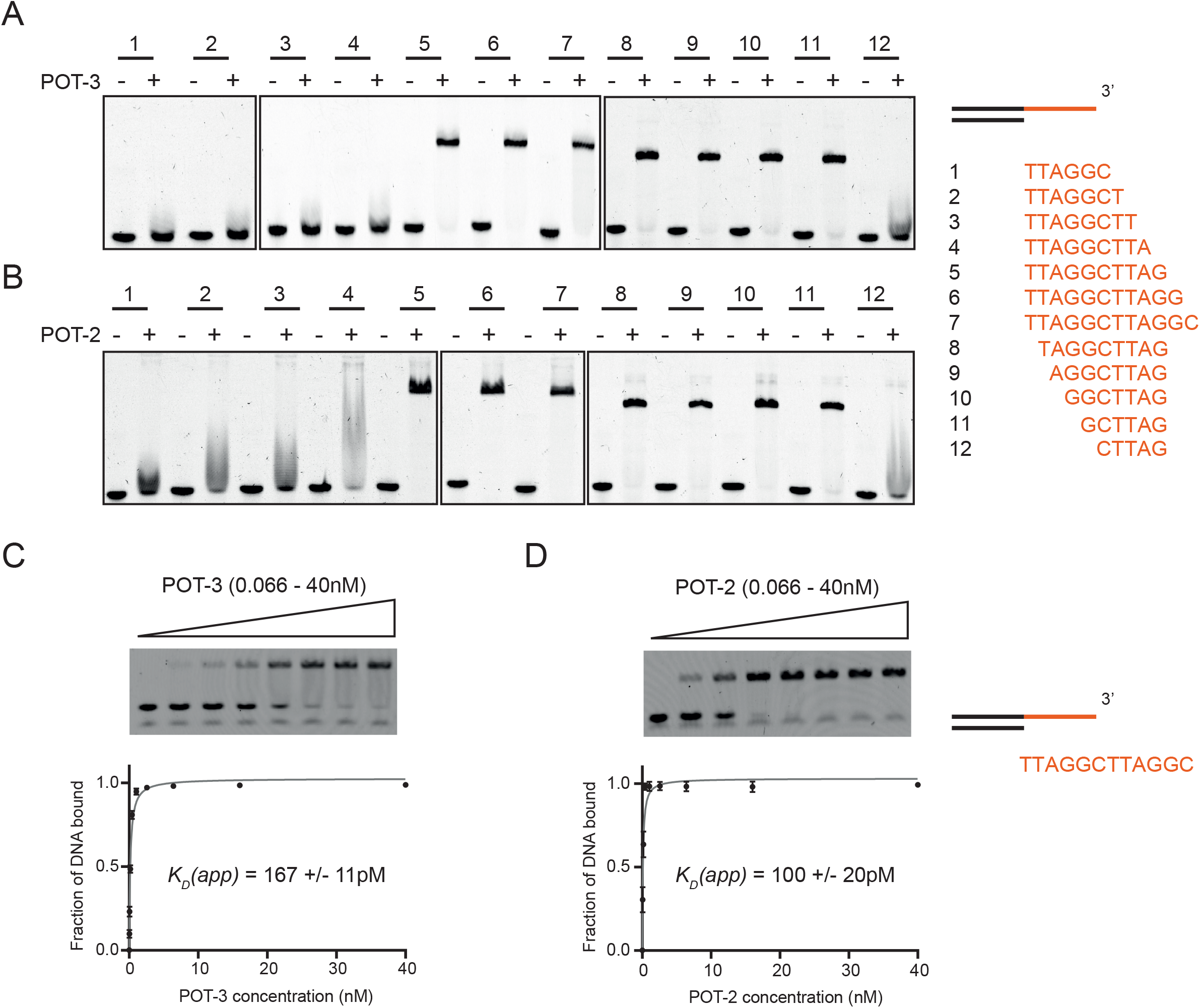
POT-3 and POT-2 bind the same minimal six-nucleotide recognition sequence, GCTTAG. **A**. 500nM POT-3 was bound to a series of DNA telomere fragments (50nM) that had 3’ overhangs increased by one nucleotide at a time from TTAGGC (fragment 1) to (TTAGGC)_2_ (fragment 7). The first fragment in this series to show binding (fragment 5) was then shortened one nucleotide at a time from the 5’ end until it was no longer able to be bound by POT-3 (fragment 12). **B**. As with part A, except POT-2 was used. POT-2 binds the same minimal sequence as POT-3 but is less selective, (compare fragment 4 in parts A and B) **C**. 0.2nM Cy5 labelled DNA containing a (TTAGGC)_2_ overhang was incubated with increasing amounts of purified POT3 (0.066-40nM). The apparent dissociation constant K_D_(app) was calculated in Prism using a one site - specific binding equation and is the mean of three replicates +/- the standard deviation. **D**. Same as in part C except using purified POT2.

POT-2 has previously been shown to also bind the telomeric G-strand [11]. We wondered whether POT-2 and POT-3 might differ in the precise sequence they bound within the G-overhang. Therefore, we carried out the same experiment as previously described to map its minimal binding motif. Surprisingly, we find that POT-2 binds to exactly the same minimal GCTTAG motif as POT-3 (Figure 3A, B). We tried estimating the binding affinities of POT-2 and POT-3 to telomeric substrates using EMSA. These revealed that both proteins have remarkably tight binding in the picomolar range. Due to the Cy5 detection limits of our scanner, we struggled to detect signal at the low DNA concentrations necessary to make accurate binding measurements. We estimate that the apparent Kds of POT-2 and POT-3 are 100pM and 160pM respectively (Figure 3C, D). Importantly, these values represent an upper limit of affinity and the true Kd of POT-2 and POT-3 for telomeric ssDNA is likely be lower than this. However, this upper limit is sufficient conclude that the affinity of POT-2 and POT-3 is still much tighter than that of human POT1, which is 9.2nM [17]. This is surprising given that the *C. elegans* POT proteins only have a single OB fold compared to the two OB-folds that human POT1 uses to bind telomeric DNA.

### POT-3 has a preference over POT-2 for binding at the 3’ end of DNA

The fact that POT-2 and POT-3 bind the same minimal motif, made us wonder how they differed. These proteins are clearly carrying out distinct functions *in vivo*. Mutation of either single gene causes a telomeric phenotype, which is not additive when both genes are deleted. This argues against both proteins performing the same role or their individual phenotype being caused by a dosage effect. We surmised that although they bound the same motif, perhaps it mattered where the motif was along the telomeric repeat.

To test this, we carried out a competition experiment where we assayed the ability of different DNA templates to outcompete pre-bound POT-2 or POT-3 complexes. One of these DNA templates, which we term the 10mer, has a TTAG<u>GCTTAG</u> overhang. Therefore, the underlined minimal binding site is at the extreme terminus of the DNA, adjacent to the 3’ hydroxyl. The second template, which we term the 12mer, has a TTA<u>GCTTAG</u>GC overhang. Here, the underlined minimal binding site is internal, two nucleotides away from the end of the DNA.

We pre-bound POT-2 to Cy5-labelled 12mer and titrated in either unlabelled 10mer or 12mer DNA. We found that there was no difference between these two competitors (Figure 4A,C). This shows that POT-2 has a similar affinity for DNA templates regardless of whether its binding site is internal or terminal. In contrast, when we repeated this with POT-3, we found that it was outcompeted more efficiently by the 10mer than by the 12mer (Figure 4B,D). This shows that POT-3 prefers to bind to sites at the ends of DNA. This end-binding preference requires proximity to the 3’ end because if the minimal binding site is switched to the 5’ end, then POT-3 is incapable of binding (Supplementary figure 1). Interestingly, POT-2 shows reduced binding but it is still capable of binding when presented with a substrate with a binding site at the 5’ end of DNA.

**Figure 4.**
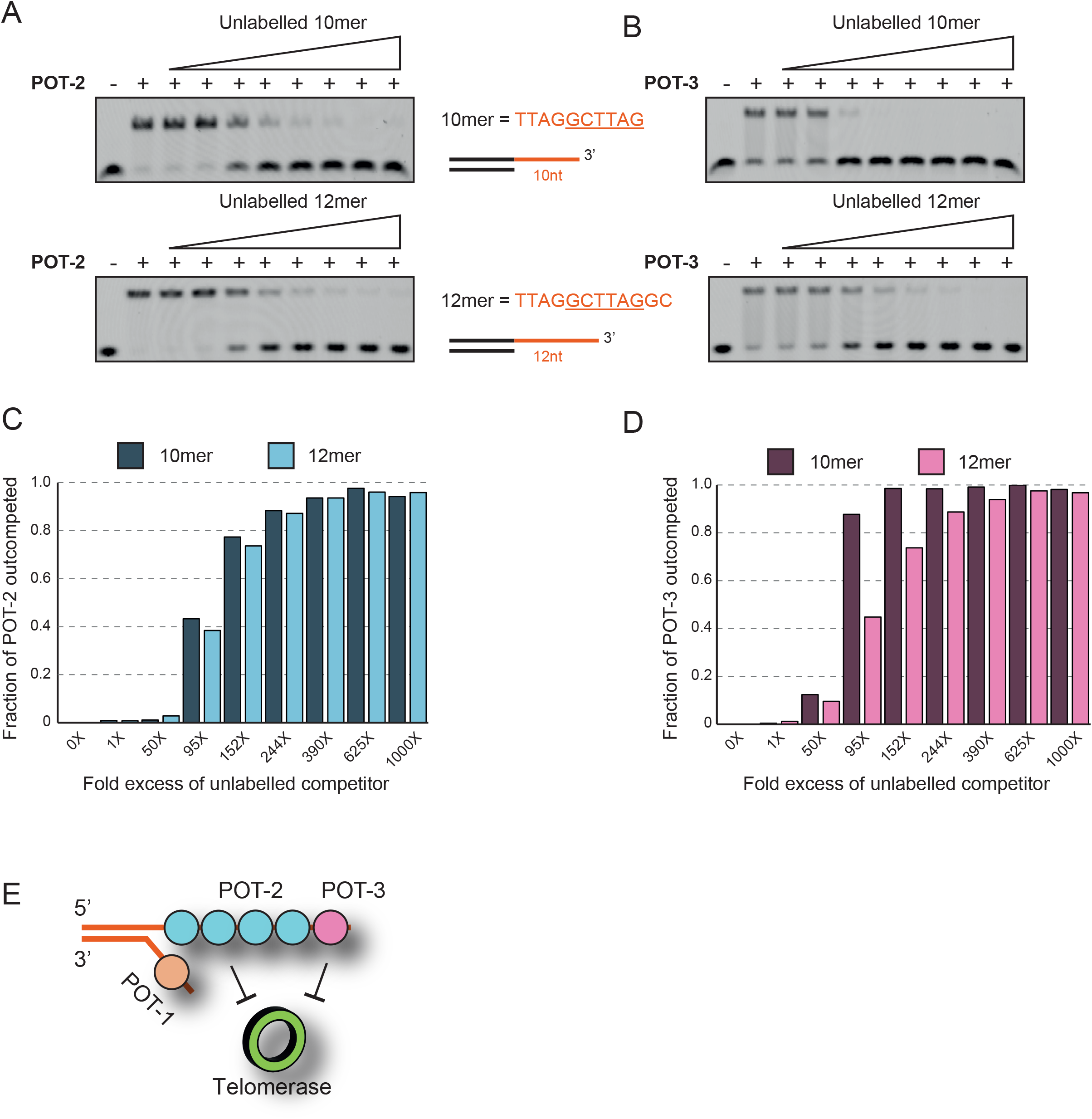
POT-3 preferentially binds its recognition sequence when it is immediately adjacent to the terminal 3’ hydroxyl. **A**. 1nM POT-2 was pre-bound to 0.2nM Cy5-labelled 12-mer DNA and incubated with increasing amounts (0.2 - 200nM) of unlabelled 10-mer or 12-mer DNA for 30mins before performing an EMSA. **B**. Same as part A except that POT-3 is used. **C** and **D**. Quantification of parts A and B plotting the fraction of POT-2 or POT-3 outcompeted by either the 10mer or 12mer DNA fragments. POT-2 can be outcompeted equally by either the 10mer or 12mer fragments, whereas POT-3 is outcompeted more efficiently by the 10mer. This indicates that it prefers its six nucleotide binding site (underlined) at the extreme 3’ end. **E**. Working model illustrating that POT-2 and POT-3 coat the telomeric G-overhang. They both repress telomerase activity to prevent the inappropriate lengthening of telomeres. However, in contrast to POT-2, POT-3 prefers to bind to the terminal repeat at the 3’ end of the G-overhang.

This increased DNA-end selectivity of POT-3 over POT-2 is also consistent with the data that POT-2 shows more promiscuous binding than POT-3 to DNA sequences that approximate its minimal binding site (substrate 4 in Figure 3B). Altogether, this shows that, although they bind the same minimal sequence, POT-3 is more selective than POT-2 and it prefers to bind the terminal telomeric repeat of the 3’ G-overhang. This region is the site of action for telomerase and exonucleases such as MRT-1, meaning that POT-3 is in a privileged location to influence telomere maintenance.

## Discussion

*C. elegans* contains three POT1-like proteins (POT-1, POT-2 and POT-3) which contain the characteristic oligonucleotide/oligosaccharide-binding (OB) fold. A fourth *C. elegans* protein (MRT-1) also contains a similar OB fold. However, in contrast to human POT1, MRT-1 additionally contains an active nuclease domain [18], suggesting a distinct function. This is supported by *mrt-1* mutants having short telomeres [18], the opposite phenotype of *pot-1* or *pot-2* mutants [11]. We show here that *pot-3(syb2415)* mutants also have long telomeres. This phenotype is different to that previously observed with *pot-3(ok1530)*. Importantly, the *ok1530* allele remains uncharacterised and there is no published evidence that it alters the POT-3 coding sequence. In, contrast, *syb2415* is a 500bp deletion that completely removes the OB fold, the only known functional domain within POT-3.

Our work and others show that all three nematode POT proteins prevent telomere elongation; however, they do so in different ways. Telomeres in *C. elegans* are unusual in that they contain 5’ C-strand overhangs (bound by POT-1) as well as 3’ G-strand overhangs (bound by POT-2 and POT-3). Thus, POT-2 and POT-3 might antagonise telomerase via direct competition. However, POT-1 likely acts via a different mechanism as it binds the opposite strand of DNA to telomerase that might involve altering its nuclear localisation [16].

We also show that *pot-2* acts epistatically with *pot-3* to prevent C-circle formation. These extra-chromosomal circles of telomeric DNA occur in a subset of cancers [13] and are thought to be generated via a combination of long telomeres and replication stress [19] as well as through inappropriate DNA repair mechanisms at telomeres [20, 21]. It has been proposed that C-circles are formed via the cleavage of a T-loop intermediate [22]. Longer telomeres may be more likely to form T-loops, which could be why *pot-2* and *pot-3* mutants have higher levels of C-circles than wildtype.

The epistasis between *pot-2* and *pot-3* in C-circle formation is interesting. We speculate that part of the function of POT-2 is to coat the bulk of the G-overhang and restrict POT-3 to the terminal repeat. Indeed, there is approximately 100x more POT-2 protein in worm embryos than POT-3 [23]. This disparity in abundance might also explain why POT-2 but not POT-3 was detected via mass spectrometry in pulldowns using the telomere binding proteins TEBP-1 and TEBP-2 [24]. Thus, in a *pot-2* mutant, POT-3 is no longer restricted to the terminal telomeric repeat and may now mislocalise to other regions such as the displaced G-strand formed next to a T-loop. We speculate that this mislocalised POT-3 makes T-loops more likely to be aberrantly processed into C-circles (Supplementary figure 2). Such a model would explain why a double *pot-2; pot-3* mutant has lower C-circle levels that a *pot-2* single mutant.

POT-2 and POT-3 bind ssDNA as monomers using a single OB-fold. This is different to most organisms where telomeric ssDNA-binding proteins often contain multiple OB folds which either contribute to DNA-binding or to protein dimerization [25]. The fact that the minimal binding site of POT-2 and POT-3 is a six-nucleotide motif means that there would be no linker DNA between adjacent POT-2/3 proteins. Indeed, we show that multiple POT proteins can fully occupy adjacent DNA binding sites (Supplementary figure 3). The lack of a DNA linker suggests that POT2/3-bound telomeric overhangs in worms are likely to be quite rigid and this in turn might antagonise T-loop and subsequent C-circle formation. The telomeric DNA of nematodes (TTAGGC) cannot form a G-quadruplex [26], as it does not contain repeated stretches of more than three sequential guanines. G-quadruplexes are often associated with genome instability but they may also play protective roles at telomeres [27]. The lack of G-quadruplexes in telomeric DNA might make their protein-bound form all the more important for telomere stability in worms.

Mutation of human POT1 causes telomere elongation [28] and is associated with cancers such as glioma [29]. The complete loss of POT1 results in DNA damage activation and telomere lengthening (but not telomere fusions) in human [30] and mouse [9] cells. Interestingly, loss of POT1 homologs in simpler eukaryotes such as moss [31] and yeasts [14, 15] have a distinct phenotype, causing increased chromosome fusions and telomere shortening instead of lengthening. Loss of either POT-1, POT-2 or POT-3 results in telomere elongation but not increased chromosome fusions [11]. This indicates that *C. elegans* POT proteins behave more like their human, rather than yeast or plant, homologs.

The high level of sequence identity between POT-2 and POT-3 suggest that they arose from a gene duplication event that underwent rapid diversification of function. This is consistent with the observation that telomeric proteins undergo particularly rapid evolution [32]. Indeed, a cursory examination of closely related *Caenorhabditis* species reveals large variability in the number of POT-like genes (data not shown). It will be important to understand how POT-1, POT-2 and POT-3 work together to bind both the telomeric C-strand and G-strand in *C. elegans* and how this influences telomere maintenance.

## Acknowledgements

We would like to thank Dr Shirley Graham for help with protein purification and Janie Olver for critical reading of the manuscript. The China Scholarship Scheme supported Xupeng Yu was supported financially by the China Scholarships Council (No.201904910787). Some strains were provided by the CGC, which is funded by NIH Office of Research Infrastructure Programs (P40 OD010440).

## Methods

### Protein purification and expression

POT2 (HFP300) or POT3 (HFP301) encoding plasmids were transformed into BL21 (DE3) pOFX34 *E. coli*. Overnight cultures were used to inoculate 2L LB media and grown at 37°C until OD600 reached 0.6, then placed at 4°C for 30 min to reduce the temperature. Cultures were induced with 0.1mM IPTG and induction was carried out at 16°C for ∼16 hours. Cells were harvested by centrifugation at 5000rpm for 10 min and frozen at -20°C until processed. The cell pellets were thawed on ice and resuspended in cold lysis buffer (500mM NaCl, 50mM Tris-HCl pH 8.0, 10mM Imidazole, 10% Glycerol) + 1x cOmpleteTM protease inhibitor tablet (Roche) and sonicated on ice. This was then centrifuged at 40,000rpm for 35 min and the supernatant loaded onto a 5mL HisTrapTM HP column (GE Healthcare). It was then wash and eluted using a gradient of Buffer A (500mM NaCl, 50mM Tris-HCl pH 8.0, 30mM Imidazole, 10% Glycerol) to Buffer B (500mM NaCl, 50mM Tris-HCl pH 8.0, 500mM Imidazole, 10% Glycerol). SDS-PAGE identified fractions containing protein of interest at highest purity. POT2 and POT3 fractions were concentrated using an Amicon® 10K concentrator at 4000rpm and the concentrated samples dialysed using SnakeskinTM 10K MWCO (Thermofisher) tubing overnight into dialysis buffer (500mM NaCl, 50mM Tris-HCl pH 8.0, 10% glycerol) at 4°C. This was then run on a HiLoad™ 16/60 Superdex 200 pg (GE Healthcare) gel filtration column equilibrated in gel filtration buffer (500mM NaCl, 20mM Tris-HCl pH 8.0, 10% Glycerol) using a flow rate of 1 mL/min. Protein concentration was determined by Nanodrop 2000 and the protein frozen in small aliquots at -80°C.

### Generation of telomeric DNA substrates

All oligonucleotides were purchased from IDT® and resuspended in autoclaved H2O to make 100μM stocks. Oligos were annealed at 1μM in ST buffer (100mM NaCl, 10mM Tris-HCl pH 8.0) in a PCR machine by heating to 95°C for 3 min and then cooling to room temperature at a rate of 1°C per minute. Annealed oligos were stored at 4°C.

### EMSA

Binding reactions were set up as follows. Appropriate concentration of protein and DNA were mixed together in binding buffer (50μg/mL BSA, 1mM MgCl_2_, 5mM DTT, 20mM Tris-HCl pH8.0, 50mM NaCl, 4% Ficoll 400). Binding reactions were incubated on ice for 10 min before loading the sample onto a 7% native polyacrylamide gel which had been pre-run for an hour. Electrophoresis was carried out in 0.5x TBE buffer for 1 hour at 100V with an ice pack to keep the temperature low.

### Strains

Unless noted otherwise, all strains were cultured at 20°C on nematode growth medium plates seeded with *Escherichia coli* OP50. A full strain list is given in Supplementary Table 2.

### Terminal restriction fragment (TRF) analysis

Worms were digested in 1x NTE buffer (100 mM NaCl, 50 mM Tris pH 7.4, 20 mM EDTA), 1% SDS and 500 μg/ml Proteinase K overnight at 65°C. Two consecutive phenol-chloroform extractions, followed by chloroform back-extraction and ethanol precipitation were carried out. DNA was eluted in 10mM Tris-EDTA (pH7.5). 5 μg purified DNA was digested overnight with HinfI and HaeIII (NEB) at 37°C and resolved on a 1% agarose gel. Following a 20-minute depurination in 250 mM HCl, the gel was washed 2x in denaturing buffer (1.5 M NaCl, 0.5 M NaOH) and 2x in neutralising buffer (1.5 M NaCl, 0.5 M Tris-HCl, pH = 8) at room temperature. DNA was transferred onto neutral nylon membrane (Hybond-NX, GE Healthcare) by capillary transfer in 10x SSC buffer (1.5M NaCl, 150 mM sodium citrate, pH 7). After briefly rinsing in 2x SSC buffer DNA was UV crosslinked at 1200 J/m^2^ and hybridised with a digoxygenin-labelled telomere probe (GCCTAA)_4_

### C-circle assay

DNA was extracted following mechanical lysis using 0.5 mm glass beads in 73 μg/ml RNase A, 9 mM EDTA and 270 mM NaCl in a cell homogeniser for 3x 20 s at 6 m/s. Proteins were denatured by adding 1% SDS and heating to 65°C for 10 minutes, followed by precipitation with a final concentration of 1.3 M Potassium Acetate pH 5.2. After additional purification using phenol:chloroform:isoamyl alcohol (15:24:1, pH = 6.7) and chloroform back-extraction, DNA was ethanol precipitated and eluted in Tris-EDTA buffer (10 mM Tris-HCl, 100 μM EDTA, pH = 7.5). This was then used in a C-circle assay as described [13]. Briefly, 0.5 μl phi29 polymerase (NEB) was added to 1ug of genomic DNA and incubated at 30°C for 8 h. This was spotted onto a neutral Hybond-N membrane, UV cross-linked (1200J/m^2^) and hybridized with a DIG-labelled (GCCTAA)_4_ probe at 37°C using DIG Easy Hyb (Roche) according to the manufacturer’s instructions.

## Notes

### Competing Interest Statement

The authors have declared no competing interest.

