## Supplementary data for "POT-3 preferentially binds the terminal DNA-repeat on the telomeric G-overhang"

### Supplementary figure 1

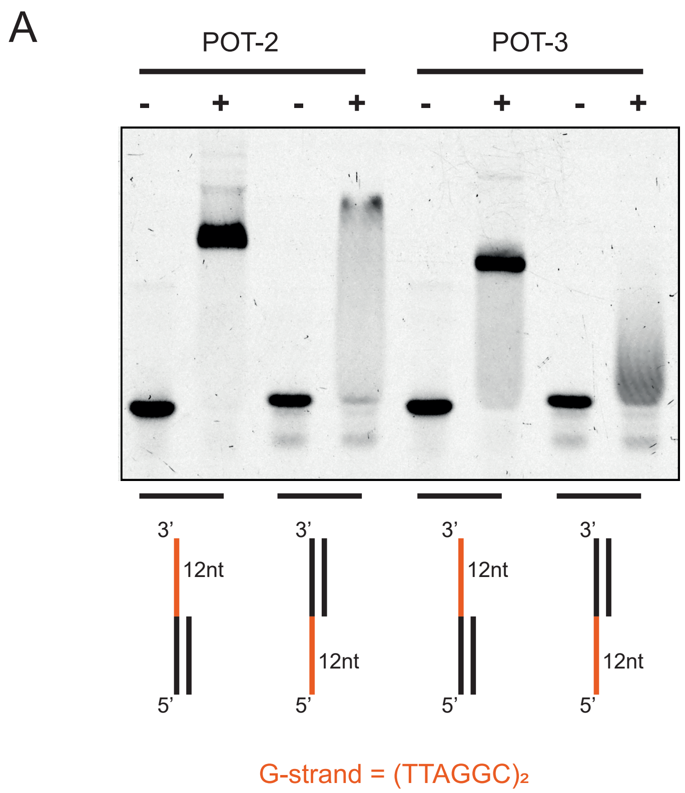

Supplementary figure 2

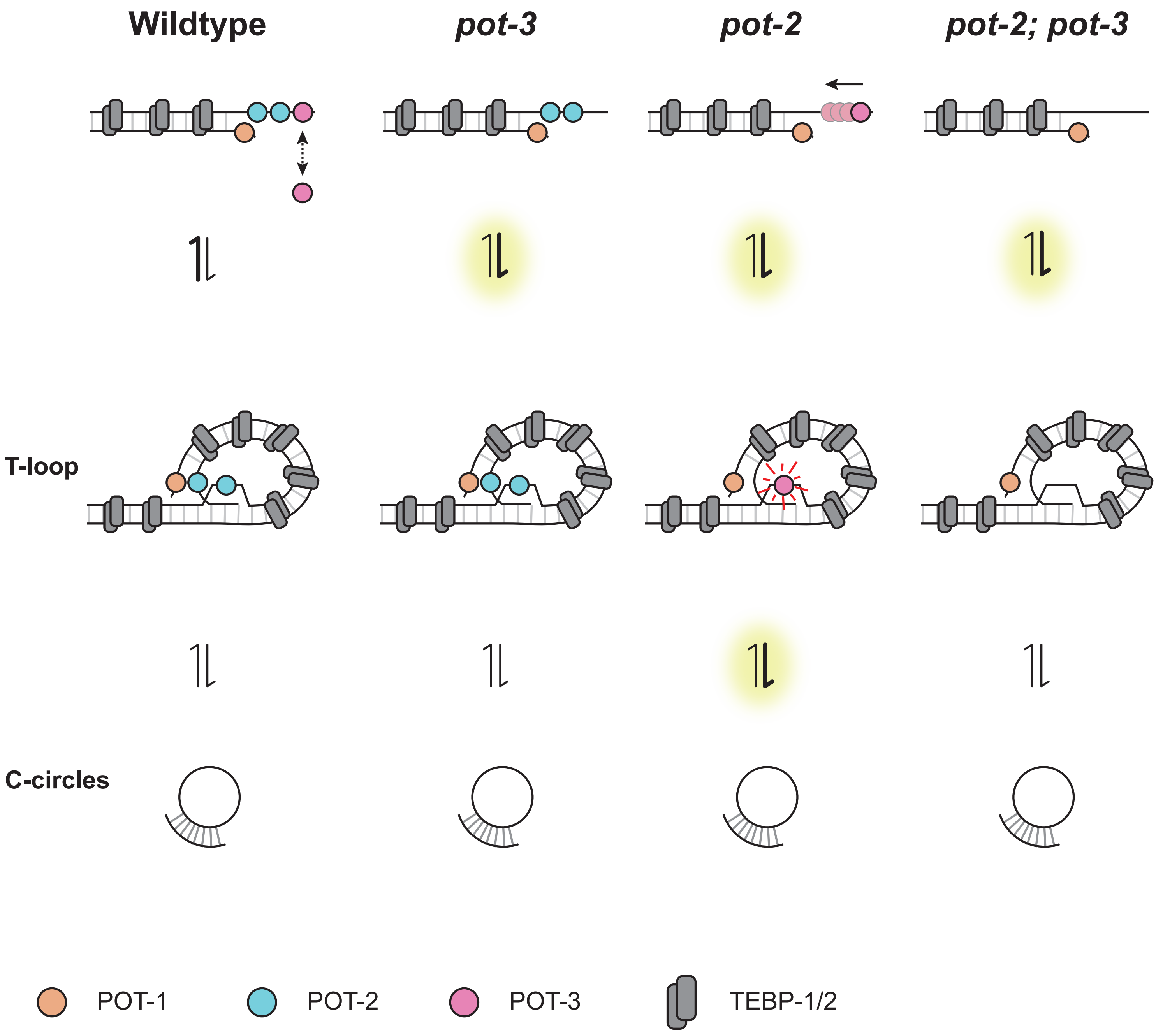

### Supplementary figure 3

| | | Elution volume | Apparent MW | Predicted MW |
|---|---|---|---|---|
| ■ (black) | POT-3 only<br>no DNA binding | 13.76ml | 27.2 KDa | 26.0 KDa |
| ■ (green) | POT-3 + $(TTAGGC)_2$<br>1 binding site | 13.25ml | 35.4 KDa | 29.8 KDa |
| ■ (red) | POT-3 + $(TTAGGC)_4$<br>3 binding sites | 11.53ml | 86.6 KDa | 85.5 KDa |

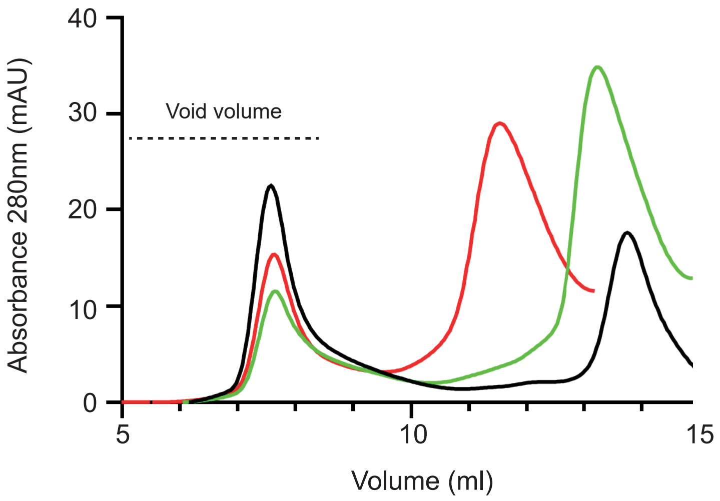

#### Supplementary table 1

##### Molecular weight ladder calibration curve (From Figure 1B)

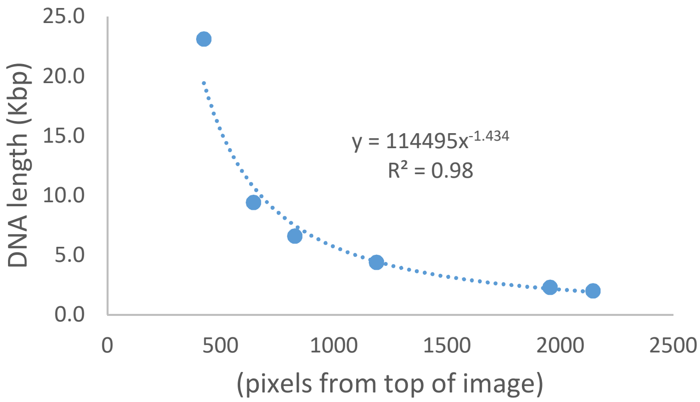


##### Telomere length

| Strain | Mode (Kbp) | Range (Kbp) |
|---|---|---|
| Wildtype | 3.9 | 2.2 - 8.6 |
| *pot-2* | 18.1 | 6.9 - 52.5 |
| *pot-3* | 9.8 | 5.5 - 23.9 |
| *pot-2; pot-3* | 13.7 | 6.3 - 21.3 |

#### Supplementary table 2

##### Strain list

| | Strain Name | Genotype | Reference |
|---|---|---|---|
| Wildtype | N2 | *Bristol strain N2* | |
| *pot-2* | HFW2 (CeOB1) | *pot-2(tm1400)* | 1 |
| *pot-3* | HFW103 | *pot-3(syb2415)* | This paper |
| *pot-2; pot-3* | HFW109 | *pot-2(tm1400); pot-3(syb2415)* | This paper |
| *trt-1* (ALT survivor) | HFW36 (c1-25) | *trt-1(ok410) unc-29(e193)* | 2 |

1. Raices, M., et al., *C. elegans* telomeres contain G-strand and C-strand overhangs that are bound by distinct proteins. Cell, 2008. 132(5): p. 745-57.

2. Cheng, C et al., *Caenorhabditis elegans* POT-2 telomere protein represses a mode of alternative lengthening of telomeres with normal telomere lengths. Proc. Natl. Acad. Sci. 109, 7805–7810 (2012).

**Supplementary Figure 1**. POT-3 requires its recognition sequence to be close to a 3' end of DNA. POT2 or POT3 (500nM) was incubated with Cy5 labelled oligonucleotide substrate (50nM) containing a non-specific duplex with TTAGGCTTAGCC overhangs at either the 3' or 5' end.

**Supplementary Figure 2**. Speculative model to explain the epistatic relationship between *pot-2* and *pot-3*. In wildtype worms, POT-3 binds the end of the G-overhang. It needs to be removed from the G overhang to reveal the terminal telomeric ssDNA that is able to invade upstream telomeric dsDNA and form a T-loop. This is unfavourable and therefore relatively few T-loops are formed. The inappropriate processing of T-loops via DNA repair pathways may facilitate the formation of C-circles. In *pot-3* worms, the end of the G-overhang is more accessible and this shifts the equilibrium (yellow highlight) to higher levels of T-loops and consequently higher levels of T-loops. In *pot-2* worms, there is not enough POT-3 to coat the G overhang and POT-3 binding is no longer restricted to 3' end. Therefore, as in *pot-3* mutants, the increased accessibility of the 3' end increases T-loop formation (yellow highlight). However, the remaining POT-3 may now bind inappropriately to the displaced G-strand within the T-loop. We speculate that this makes the T-loop more recombinogenic, shifting the equilibrium (yellow highlight) towards the higher levels of C-circles. This increase in C-circle levels in *pot-2* can be suppressed by removing POT-3, such that a *pot-2, pot-3* double mutant now has lower C-circle levels than in the single *pot-2* mutant and instead now shows similar C-circle levels to a single *pot-3* mutant.

**Supplementary Figure 3**. Analytical gel filtration reveals that POT3 binds telomeric sequences as monomers. Purified POT-3 was incubated with either no DNA, $(TTAGGC)_2$ or $(TTAGGC)_4$ ssDNA and run on a Superose® 12 10/300 GL column. The chromatogram does not show peaks of DNA absorbance alone, which were removed for simplicity.
